# The ebb and flow of heteroplasmy during intra-species hybridization in *Caenorhabditis briggsae*

**DOI:** 10.1101/623207

**Authors:** Shadi Adineh, Joseph A. Ross

## Abstract

Mitochondria are typically maternally inherited. In many species, this transmission pattern is produced by sperm-borne mitochondria being eliminated either from sperm before fertilization or from the embryo after fertilization. In the nematode *Caenorhabditis briggsae*, repeatedly backcrossing hybrids to genetically diverse males can elicit paternal mitochondrial transmission. Studies of other taxa also suggest that hybridization increases paternal mitochondrial transmission. Thus, hybrid genotypes might disrupt the systems that normally prevent paternal mitochondrial transmission. Given the reliance of a number of genetic analyses on the assumption of purely maternal mitochondrial inheritance, it would be broadly valuable to learn more about the processes embryos employ to prevent sperm-borne mitochondria from persisting in offspring, as well as the circumstances under which paternal transmission might be expected to occur. To quantify the tempo of paternal mitochondrial transmission in hybrids, we assessed the presence of paternal mitotypes in replicate lines at three timepoints spanning fifteen generations. All lines exhibited paternal mitochondrial transmission. However, this heteroplasmy always then resolved to homoplasmy for the maternal mitotype. Additionally, one nuclear locus exhibited allele transmission ratio distortion that might reflect mito-nuclear co-evolution. This study frames the genetic architecture of a hybrid genetic incompatibility that leads to paternal mitochondrial transmission and to a reduction in hybrid fitness.

## INTRODUCTION

Mitochondria play a pivotal role in cellular metabolism (Taylor and Turnbull 2005). Much of the mitochondrial genome (mtDNA) encodes genes essential for transcription and translation as well as the oxidative phosphorylation system (Bereiter-Hahn and Voth 1994; Timmis, et al. 2004). Nuclear genes also encode protein subunits of electron transport chain complexes. The reliance of mitochondrial function on genes encoded by two genomes established an evolutionary dynamic where mitochondrial and nuclear genes co-evolve (Blier, et al. 2001). Deleterious mutations in the mitochondrial genome can be compensated by mutations in nuclear genes encoding adjacent protein subunits (Rand, et al. 2004). When hybridization occurs between lineages carrying different co-evolved genomes, hybrid breakdown can result from separation of co-evolved genes and subsequent mitochondrial dysfunction (Gershoni, et al. 2009).

Eukaryotes typically exhibit inheritance of mtDNA from their mothers (uniparental inheritance) (Gyllensten, et al. 1991; Hurst 1992; Fontaine, et al. 2007), although empirical data have occasionally shown the presence of both paternal and maternal mtDNA haplotypes (mitotypes) in one individual (e.g. (Birky 1995)). In some species, like mussels, uniparental inheritance does not occur (Zouros, et al. 1992; Zouros, et al. 1994). The existence of predominantly (if not exclusively) maternal mtDNA inheritance might prevent inheritance of sperm mtDNA that carry deleterious mutations (Mazat, et al. 2001). The presence of two mitotypes (paternal and maternal) within a single zygote (termed heteroplasmy) might also be unfavorable (Hurst 1992; Sharpley, et al. 2012) (Sato and Sato 2013), but see also (Ye, et al. 2014). Thus, maternal mitochondrial inheritance might exist to ensure homoplasmy, the presence of a single mitochondrial haplotype in a cell.

At least two paternal mitochondrial elimination mechanisms exist: mitochondria in sperm can be destroyed prior to fertilization, and mitochondria that enter the oocyte can be eliminated after fertilization. For example, mammalian sperm enter the oocyte at fertilization, and their mitochondria are tagged with ubiquitin for destruction by the proteasome (Sutovsky, et al. 1999); mouse mtDNA are also eliminated prior to fertilization (Luo, et al. 2013). In the nematode *C. elegans*, evidence suggests roles for ubiquitin and mitophagy in removing paternally-transmitted mitochondrial following fertilization (Al Rawi, et al. 2011; Sato and Sato 2011; Zhou, et al. 2016). Such evidence of active elimination of paternal mitochondria in the embryo suggests that a molecular mechanism(s) in the oocyte detects some signal identifying sperm-borne mitochondria.

Here, we have investigated the dynamics of paternal mitochondrial transmission (PMT) in intra-specific *C. briggsae* hybrids, which are known to sustain occasional PMT (Chang, et al. 2016; Ross, et al. 2016). We created reciprocal sets of triplicate cytoplasmic-nuclear hybrid (cybrid) lines initiated by mating the genetically distinct *C. briggsae* wild isolate strains AF16 and HK104, which are about one percent divergent in the nuclear genome (Hillier, et al. 2007). These cybrid lines experimentally combine the nuclear genome of one strain with the mitochondrial genome of the other. We genotyped individuals at various generations during line production to assess the presence of paternal mitochondria and found that all lines exhibited PMT during line production, but heteroplasmy eventually resolved in favor of homoplasmy for the maternal mitotype in all lines. By genotyping genetic markers in the nuclear genome, we also discovered a pattern of mito-nuclear linkage disequilibrium that implies the presence of co-evolved loci. Together, these results support the interpretation that an inter-genomic genetic incompatibility precipitates PMT in hybrids.

## MATERIALS AND METHODS

AF16 and HK104 wild isolates were obtained from the Caenorhabditis Genetics Center, which is funded by NIH Office of Research Infrastructure Programs (P40 OD010440). To best control for the effect of hybridization in this experiment, the reciprocal control lines (FV83–5, FV62–4) and the cybrid lines (FV59–61, FV86–8) were all initiated from the F1 hybrid generation, with serial P0 paternal backcrosses then used to create cybrid lines and maternal backcrosses used to create control lines (Figure 1). Strain husbandry and cybrid crosses were conducted as previously described (Chang, et al. 2016).

**Figure 1.**
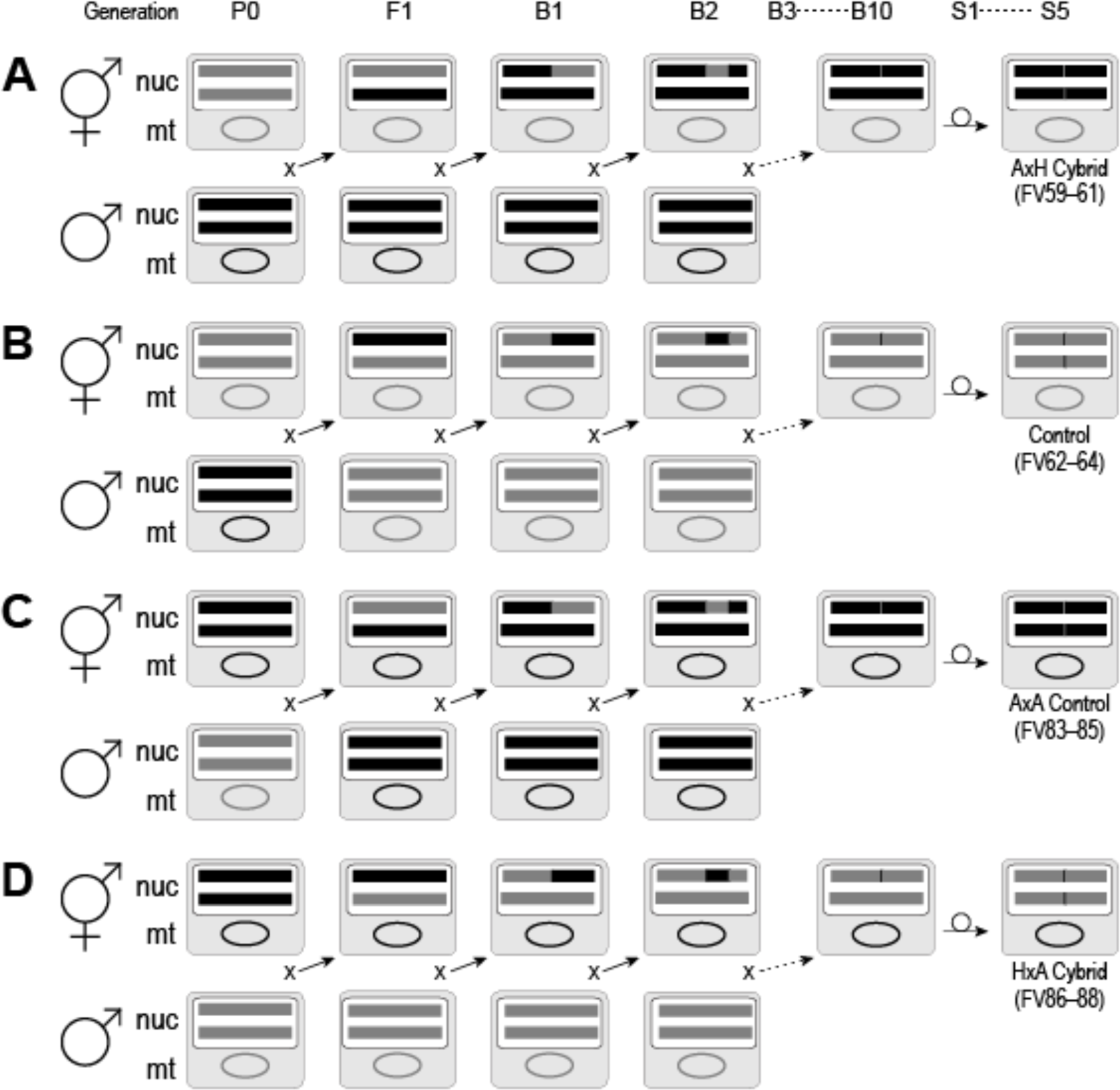
Cross design and expected genotypes of lines. The average neutral expectation allele retention patterns in hybrid lines are depicted using gray shading for HK104 alleles and black for AF16 alleles. An example diploid autosome pair (“nuc,” horizontal lines) and mitochondrial genome (“mt,” circular genome) are depicted for the following generations: P0, F1, first and second backcross generations (B1, B2), B10, and the fifth generation of serial selfing (S5). Control crosses in panel B (FV62–64) and in panel C (FV84–86), in which an F1 hybrid is serially backcrossed to males from the P0 maternal strain, reconstitute the P0 maternal nuclear genotype on the same cytoplasmic background. Experimental crosses in panel A (FV59–61) and in panel D (FV86–88), in which an F1 hybrid is serially backcrossed to the P0 paternal strain, produce cytoplasmic-nuclear hybrid (cybrid) lines that reconstitute the P0 paternal nuclear genotype on the maternal cytoplasmic background.

Primer sequences for amplified fragment length polymorphism (AFLP) genotyping of nuclear loci (II: cb-m26, III: cb-m205, IV: cb-m172 and cb-m67, V: cb-m103, X: cb-m124 and cb-m127) and for restriction fragment length polymorphism (RFLP) genotyping of mtDNA loci (cb18178, cb34440) were obtained from (Koboldt, et al. 2010).

Single-worm PCR involved isolating an individual adult hermaphrodite nematode in 5 µL of single-worm lysis buffer (50 mM KCl, 10 mM TRIS pH 8.3, 2.5 mM MgCl2, 0.45% nonidet P40, 0.45% tween-20, 0.01% gelatin, 1 mg/mL proteinase K). After snap-freezing at −80°C, each worm was incubated at 60°C for 60 m. and then at 95°C for 15 m. 1 µL of worm lysate was then used as template DNA for AFLP and RFLP PCR and analysis as described (Chang, et al. 2016).

The nematode lines will be made available to qualified researchers upon request.

## RESULTS AND DISCUSSION

We speculated that negative mito-nuclear epistatic interactions in cybrids might elicit PMT in the experimental lines. The presence of a paternal mitotype in any hybrid generation would reveal PMT, and we expected that nuclear allele frequencies would not deviate from the values expected for selectively neutral marker loci. We genotyped eight individuals from each of the twelve lines at two nuclear loci and one mitochondrial locus. Limited locus sampling was necessary given the small amount of genomic DNA obtained from a single worm.

The HK104 allele frequencies of each line at the B5, B10 and S5 generations at two nuclear loci and one in the mtDNA are plotted in Figure 2. Each data point represents the frequency calculated from the successful genotyping of between two to eight individuals. Panels A and B of Figure 2 display the change in the frequency of the HK104 alleles at nuclear loci and in the mtDNA in reciprocal crosses designed to create cybrids. The expected HK104 nuclear allele fraction values at each generation (solid line, open diamond) are based on the neutral expectation of allele inheritance. HK104 allele fraction values were quite variable at the B5 and B10 generations. Some of this variation is likely due to the small number of individuals (between two and eight) successfully genotyped at these generations; this variation in replication was caused by the vagaries of success of single-worm genotyping by PCR. The expected HK104 mitochondrial allele fraction values at each generation (solid line, open circle) are based on the assumption of strictly maternal inheritance (Figure 2). Thus, in AF16xHK104 crosses, at every generation, we expected that all individuals would contain only HK104 mtDNA (HK104 allele fraction = 1, Figure 2A). In the reciprocal cross, all individuals should only contain AF16 mitotypes (HK104 allele fraction = 0, Figure 2B). If, at any point, the observed mitochondrial allele fraction value deviates from 1 or 0, respectively, then that would indicate that at least one individual genotyped in that generation contained paternal mtDNA. In the B5 and B10 generations of both cybrid crosses, we did observe paternal mitotypes. For example, in the B10 generation of the AF16xHK104 cybrid cross (Figure 2A), four of eight individuals in strain FV59 had paternal mitotypes; five of eight FV60 individuals had paternal mitotypes (Figure 2B). Interestingly, we also observed evidence of paternal mitotypes in the B5 and B10 generations in the control crosses (Figures 2C, D).

**Figure 2.**
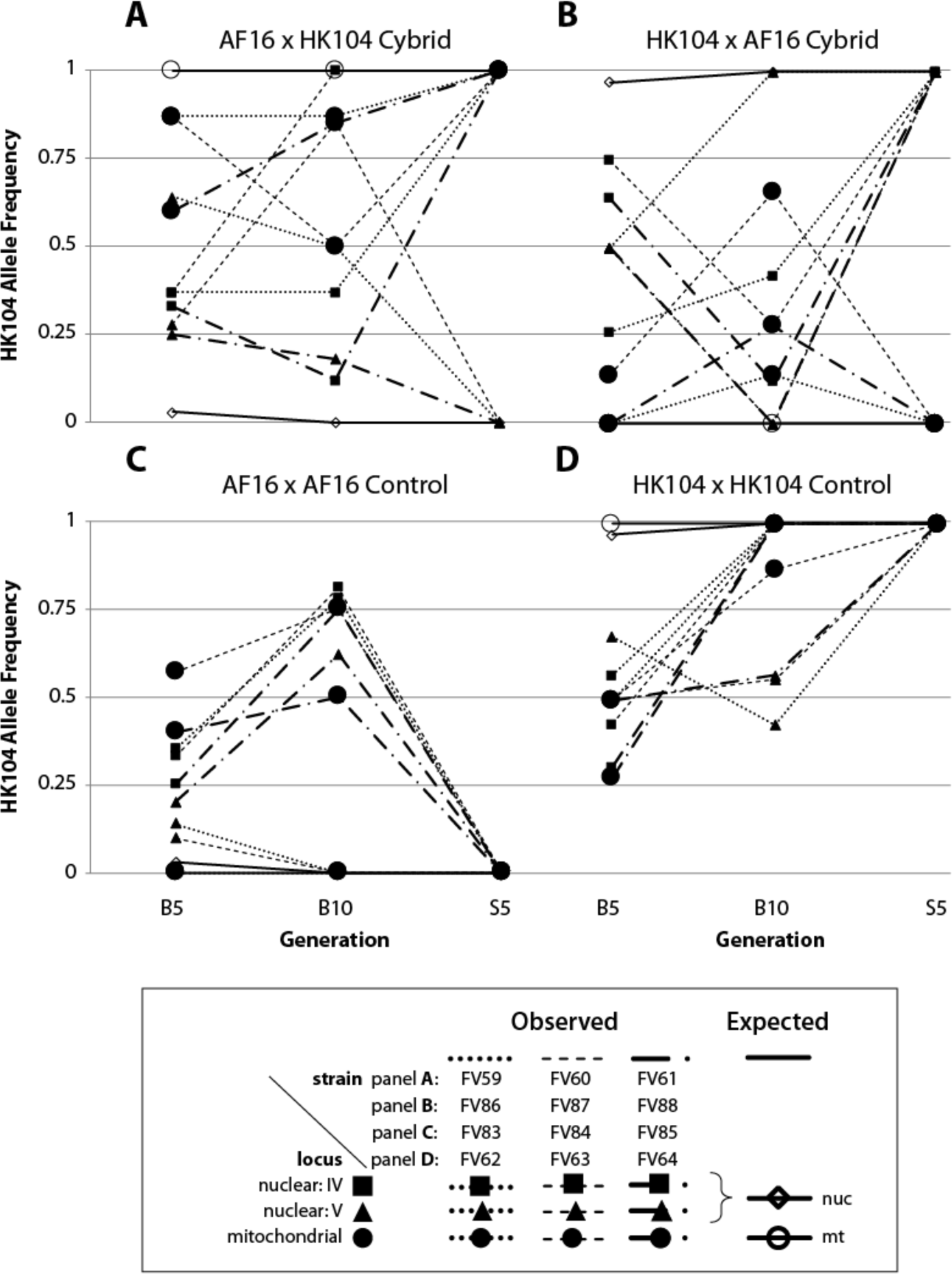
HK104 allele fraction values in cybrid and control lines. The HK104 allele frequencies at two nuclear loci (cb-m67(IV) and cb-m103(V)) and one mtDNA locus are shown for generations B5, B10 and S5. Panels A and B contain genotype data for two reciprocal crosses that produced three replicate cybrid lines each; panels C and D show data from triplicate reciprocal control lines. Panel letters correspond to the cross design panel lettering in Figure 1. Each panel contains eleven data series. Nine comprise three loci (distinguished by marker shapes (squares (IV), triangles (V), and circles (mtDNA))) in each of three replicate lines (distinguished by dotted, dashed, and dotted-dashed lines). The two expected allele frequency patterns for nuclear (diamond) and mitochondrial (circle) loci are plotted as solid lines.

Once each line reached the fifth generation of serial selfing (S5), at which homozygosity throughout the nuclear genome was expected (Figure 1), genomic DNA from populations of >10,000 individuals was extracted from all twelve lines. Each sample was genotyped at seven loci on four nuclear chromosomes and two on the mtDNA to confirm the expected genotypes. On chromosomes II, V and X, the genotypes agreed with expectations based on the cross design: all lines contained only paternal alleles at the loci we genotyped (Figure 3). In these lines, mitochondrial genotypes also concurred at this timepoint: maternal mtDNA were exclusively detected (Figure 3). Only chromosome IV loci demonstrated a different allele retention pattern. At a nuclear locus, each line has an average 97% probability of retaining only paternal alleles after five generations of selfing under the neutral expectation. However, in both triplicate sets of cybrids, one of two chromosome IV loci (cb-m172) exclusively retained the P0 maternal allele, while the six control lines exclusively retained the P0 paternal allele (Figure 3). At a second chromosome IV locus (cb-m67), the paternal allele was retained in all three triplicate cybrids for one cross direction and not the other (Figure 3). This bias toward retention of maternal nuclear alleles is consistent with the possibility that a chromosome IV allele more tightly linked to cb-m172 than to cb-m67 has co-evolved with the mtDNA. Such an epistatic interaction might ensure that the only individuals surviving line creation were those that inherited the HK104 cb-m172 allele with the HK104 mitotype and those that inherited the AF16 cb-m172 allele with the AF16 mitotype. This interpretation of positive mito-nuclear epistasis is supported by evidence of mito-nuclear co-evolution between chromosome IV alleles and mitotype in AF16-HK104 advanced-intercross recombinant inbred lines (Ross, et al. 2011).

**Figure 3.**
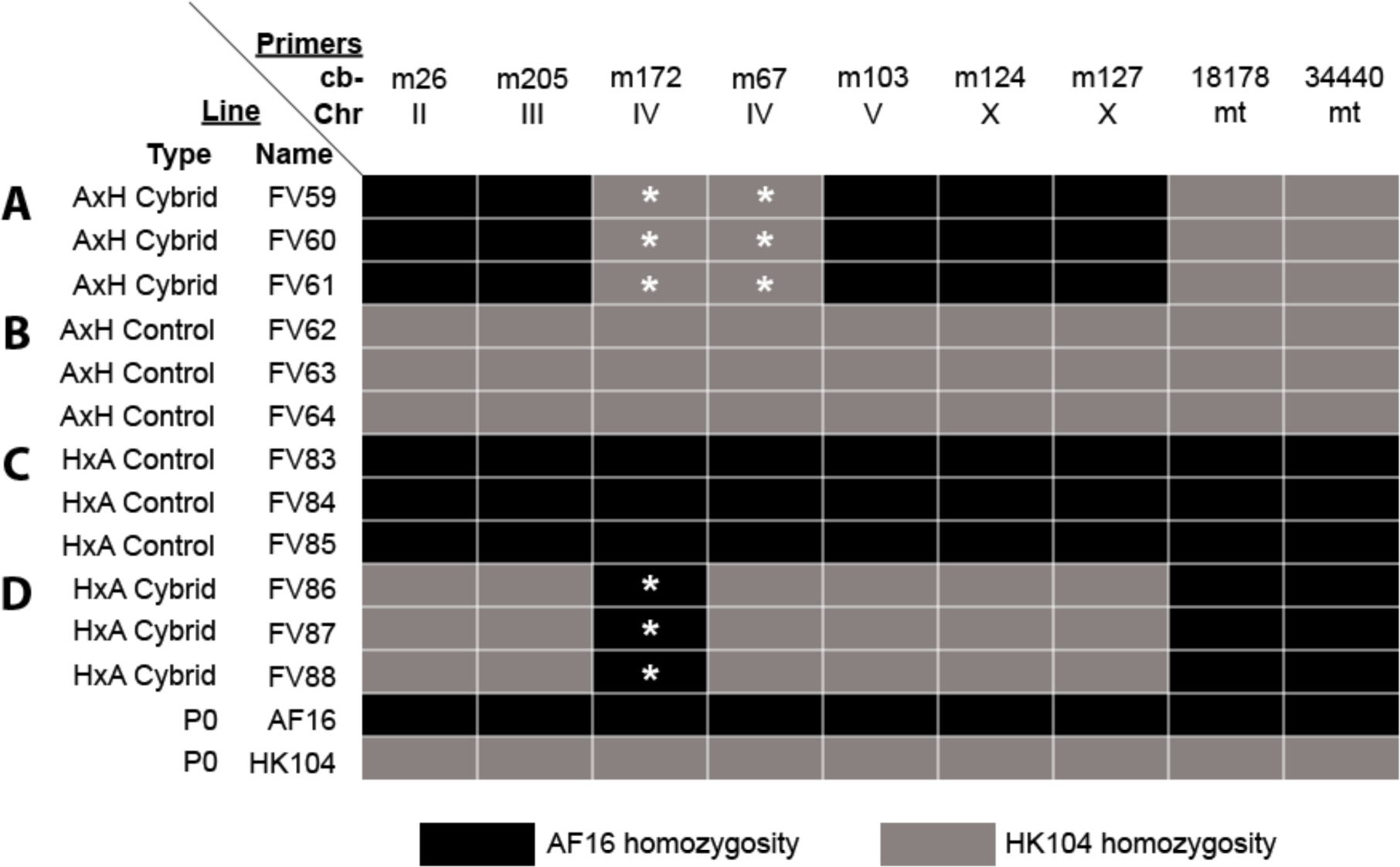
S5 generation line genotypes. Following cybrid and control line hybridization, each line (FV##) was propagated by selfing for five generations to generate homozygosity across the nuclear genome (Figure 1). DNA extracted from a population of individuals from each line was genotyped at nuclear and mitochondrial AFLP and RFLP markers (“cb-”) on multiple chromosomes (“Chr”). Control genotypes were also obtained from the P0 parental strains AF16 and HK104. Panel letters correspond to the cross design lettering in Figure 1. Panels A and D contain the genotypes of the replicate cybrid lines; panels B and C contain the control line genotypes. Genotype data are depicted by shading (black for AF16 homozygosity; gray for HK104 homozygosity). White asterisks indicate genotypes that do not agree with the expectation from the cross design.

Attempts to quantify PMT frequencies in natural populations have, in at least one case, revealed no paternal mitotypes in offspring (Wolff, et al. 2008). In *Drosophila melanogaster*, heteroplasmy is common in nature and can take many generations to resolve to homoplasmy (Nunes, et al. 2013). However, in hybrids, studies from multiple taxa suggest that PMT increases in frequency. For example, a study of the bird *Parus major* showed that introgression of a paternal mitochondrial haplotype occurred following intra-species hybridization and generated a single heteroplasmic individual (Kvist, et al. 2003). Heterospecific and conspecific hybridization in mice, fruit flies, chickens, and other nematodes have also produced evidence of PMT (Shitara, et al. 1998; Sherengul, et al. 2006; Hoolahan, et al. 2011; DeLuca and O’Farrell 2012; Wolff, et al. 2013; Alexander, et al. 2015). Thus, a growing body of empirical evidence suggests that hybridization increases the frequency of PMT. We favor the interpretation that hybridization might separate co-evolved alleles involved in maternal detection and elimination of sperm-transmitted mitochondria.

The current study adds to prior evidence of PMT in intra-species *C. briggsae* cybrids (Chang, et al. 2016; Ross, et al. 2016). However, other AF16-HK104 hybrids, like a panel of 167 advanced-intercross recombinant inbred lines, do not contain paternal mitotypes (Ross, et al. 2011). This difference suggests a genetic architecture responsible for PMT that is specific to genotypes generated by paternal backcrossing (e.g. mito-nuclear epistasis) and not related simply to having a particular hybrid nuclear genotype. The possibility of a complex genetic basis of PMT has been noted (Dokianakis and Ladoukakis 2014). It is likely that our findings are not universal, because backcrosses in mice have revealed a lesser extent of PMT (Gyllensten, et al. 1991). Paternal backcross designs in other taxa have found no evidence of PMT (Lansman, et al. 1983; Avise and Vrijenhoek 1987), although it has been suggested that, in Drosophila, such repeated hybridization is important for PMT (Wolff, et al. 2013).

We observed paternal mitotypes following hybridization in all lines. Why might we have observed paternal mitotypes even in our control strains? It is possible that, because our experimental and control strains were all initiated from F1 hybrids, they all had an opportunity to experience PMT before the cross designs diverged to P0 paternal backcrosses (to generate experimental cybrid lines) and P0 maternal backcrosses (to generate control lines).

Despite low replication of genotype collection in the B5 and B10 generations, three salient data trends are evident. First, in each of the twelve lines, paternal mitotypes were always detected in at least one individual. Second, the S5 generation lines all revealed consistent and homozygous genotypes at each locus across all three replicates. Third, only loci on chromosome IV at S5 ever contradicted the expected genotype based on the cross design. In sum, the evidence of PMT into offspring following hybridization in an intra-species cross supports previous observations in *C. briggsae* (Chang, et al. 2016; Ross, et al. 2016) as well as in other taxa. Further, the nuclear allele retention pattern observed on chromosome IV might be evidence of mito-nuclear epistasis that, when disrupted at hybridization, causes fitness reduction. Future studies of this *C. briggsae* hybrid system will be valuable to elucidate the genetic interactions that facilitate the elimination of paternal mitotypes at fertilization and further challenge the dogma of strictly maternal mitochondrial inheritance (Carelli 2015).

## ACKNOWLEDGMENTS

We thank Joel Rodriguez and Chris Jorgensen for experimental support. This work was supported by National Institute of General Medical Sciences at the National Institutes of Health (SC2-GM113727 to J.A.R.) and a Faculty-Sponsored Student Research Award from the College of Science and Mathematics, CSU Fresno.

